# *Cis*-regulatory fragments from the *dissatisfaction* gene identify novel mating behavior neurons in female *Drosophila*

**DOI:** 10.1101/2025.08.18.670925

**Authors:** Julia A. Diamandi, Kara E. Miller, Troy R. Shirangi

**Affiliations:** Villanova University, Department of Biology, 800 East Lancaster Ave, Villanova, PA 19085

## Abstract

During *Drosophila* courtship, males chase and sing to females, while females perform abdominal behaviors to indicate their willingness to mate. The nerve cord circuits in females that produce their abdominal behaviors are poorly characterized. We recently identified an anatomically diverse population of abdominal interneurons, the DDAG neurons, which express the *Tlx*-like nuclear receptor *dissatisfaction* (*dsf*) and influence several female mating behaviors. Here, we searched the *dsf* locus for *cis*-regulatory enhancer fragments that regulate its spatial expression in the adult and larval central nervous system. We found several enhancers, most located within two introns, that drove reporter expression in subsets of *dsf*-expressing neurons throughout the brain and nerve cord. Using one of these enhancers, we genetically isolated a single subtype of female-specific DDAG local interneurons. Optogenetic activation of these neurons triggered vaginal plate opening in both unmated and mated females, a behavior used by *Drosophila* females to signal receptivity to courting males. Our findings offer new reagents to target *dsf*-expressing cells and new insights into the neural substrates in *Drosophila* females that express their mating decisions during courtship.

## Introduction

Females across animal clades often perform intricate motor behaviors to signal their mating decisions to courting males. In *Drosophila*, females respond to a male’s courtship song with subtle abdominal behaviors that convey either acceptance or rejection, depending on their mating status (Aranha and Vasconcelos 2018). Unmated females are generally receptive and indicate their willingness to mate by slowing down, spreading their vaginal plates, exposing their ovipositor and permitting copulation (Mezzera et al. 2020; Wang et al. 2021). After mating, *Drosophila* females become unreceptive for several days, until they run out of sperm. During this period, mated females still open their vaginal plates in response to male courtship, but subsequently fully extrude their ovipositor, a behavior that signals rejection (Mezzera et al. 2020; Wang et al. 2020).

In both female and male flies, courtship behaviors are largely innate and emerge during metamorphosis through the actions of two key sexual differentiation genes, *doublesex* (*dsx*) and *fruitless* (*fru*) (Oren-Suissa and Shirangi 2025). These genes have been instrumental in uncovering the neural circuits governing courtship behaviors in both sexes. The circuitry in the ventral nerve cord that controls male courtship song, for example, has been elucidated in remarkable detail (Lillvis et al. 2024). In contrast, the neural circuits driving the abdominal motor behaviors of females during courtship remain less well understood.

We recently identified a small, sexually dimorphic population of interneurons in the adult abdominal ganglion—termed the DDAG neurons—that co-express *dsx* and the *Tlx*/*tailless*-like nuclear receptor *dissatisfaction* (*dsf*) (Duckhorn et al. 2022). These neurons are anatomically diverse, comprising at least five subtypes that influence various female mating behaviors, including vaginal plate opening in unmated females and ovipositor extrusion and egg-laying in mated females (Duckhorn et al. 2022; Diamandi et al. 2024). In previous work, we found that two of these subtypes, DDAG_C and DDAG_D, are specifically involved in ovipositor extrusion (Diamandi et al. 2024). The behavioral contributions of the remaining DDAG neurons, however, remain unclear.

In this study, we searched the *dsf* locus for enhancer elements that drive reporter expression in subsets of *dsf*-expressing neurons in both the larval and adult central nervous system. We identified one enhancer fragment that enabled genetic access specifically to the DDAG_B neurons. Optogenetic activation of these neurons triggered vaginal plate opening in females regardless of their mating status, suggesting that DDAG_B neurons are part of the motor circuitry controlling this behavior. However, silencing DDAG_B neurons had no detectable effect on female courtship behaviors, implying that other circuit elements may compensate for their loss. Our findings provide new genetic tools for accessing *dsf*-expressing neurons and offer novel insights into the abdominal circuits that regulate female mating behaviors.

## Methods and Materials

### Fly Stocks

*Drosophila melanogaster* stocks were maintained on standard cornmeal and molasses food at 25ºC and 50% humidity in a 12-hr light/dark cycle unless otherwise noted. Fly stocks used in this study were: *Canton S, dsf*^Gal4^/CyO (Duckhorn et al. 2022), *dsf*^p65AD::Zp^/CyO (Diamandi et al. 2024), *pJFRC79-8XLexAop2-FlpL* (attP40), *pJFRC41-10XUAS-FRT-STOP-FRT-myr::gfp* (su(Hw)attP1), *pJFRC56-10XUAS-FRT-STOP-FRT-kir2*.*1::gfp* (attP2), *20XUAS-FRT-STOP-FRT-CsChrimson::mVenus* (VK5), and *VT026005-Zp::GDBD* (attP2).

### Construction of *Dsf_CRE* transgenes

All regulatory fragments from *dsf* were cloned into pENTR/D and then Gateway cloned into pBPnlsLexA::p65Uw (Addgene Plasmid number: 26230). All *Dsf_CRE-LexA::p65* transgenes were inserted into the attP40 landing site. Forward and reverse primers used to amplify each fragment were the following:

Dsf_CRE_1:

5’-CACCTAACTCAATTCCCCAATTCTATCCAAGG-3’

5’-TGACGGACCAACTGCGAATGAAAC-3’

Dsf_CRE_2:

5’-CACCTCATTTGCTGTAATCGCATCAGGCC-3’

5’-CCTTGCAGGGAATGTCCAACAGGC-3’

Dsf_CRE_3:

5’-CACCAGCGAATCGCGTCGGCATCTGAAC-3’

5’-CCTTGCAGGGAATGTCCAACAGGC-3’

Dsf_CRE_4:

5’-CACCAACACAGGGGAATGCTCTATTCAAC-3’

5’-TTGACAAGTGCCGAGTGTTGAACTTA-3’

Dsf_CRE_5:

5’-CACCAACACAGGGGAATGCTCTATTCAAC-3’

5’-TTCCAGTCGGCCATGAACAAGGATG-3’

Dsf_CRE_6:

5’-CACCTCTCGCCGTGGGAACTTGCCAGCTG-3’

5’-ATCAGCATGCCACCGTACTTTAGAGC-3’

Dsf_CRE_7:

5’-CACCTGCAACTTGGGTTTCCTAGGACCGC-3’

5’-AGGAGTCAAGCAGCAAATCGATGG-3’

### *in situ* Hybridization Chain Reaction

Nervous systems were dissected in 1X phosphate-buffered saline (PBS) and fixed in 4% paraformaldehyde in PBS for 35 minutes at room temperature, then rinsed three times in PBS with 1% Triton X-100 (PBT) and washed in PBT for 20 minutes. The HCR™ Gold RNA-FISH Kit (Molecular Instruments) was used to perform *in situ* hybridization chain reaction as previously described (Duckhorn et al. 2022). HiFi Probe Hybridization Buffer was pre-warmed to 37 °C for 15–20 minutes before use. Tissues were rinsed 1–2 times in pre-warmed HiFi Probe Hybridization Buffer, then incubated for 30 minutes at 37°C in fresh buffer. 1 µL of HCR™ HiFi Probe (GFP X1) was added to 250 µL Probe Hybridization Buffer, and tissues were incubated overnight (∼16 hours) at 37 °C. The following day, tissues were washed four times for 15 minutes at 37°C in pre-warmed 1X HiFi Probe Wash Buffer and three times for 5 minutes in 5X SSCT (UltraPure 20X SSC Buffer, Invitrogen #15557044, diluted in water) at room temperature. Samples were then pre-amplified in Gold Amplifier Buffer for 30 minutes at room temperature. 4 µL of hairpins (X2 647 HCR™ Gold Amplifier) were snap cooled for 30 minutes in the dark before being adding 100 µL of Amplifier Buffer to make the final Hairpin solution. Tissues were incubated in the Hairpin solution overnight at room temperature, protected from light. On the third day, tissues were washed twice for 5 minutes, twice for 30 minutes, and once for 5 minutes in 5X SSCT at room temperature, then rinsed three times in PBT and washed for 20 minutes. Samples were mounted in VectaShield (Vector Laboratories, Cat. #H-1000-10) and imaged on a Leica TCS SP8 Confocal Microscope at 40X magnification.

### Immunohistochemistry

Immunohistochemistry was performed as previously described (Duckhorn et al. 2022). Briefly, nervous systems were dissected in 1X PBS, fixed in 4% paraformaldehyde in PBS for 35 minutes, rinsed and washed in PBT. We did not block. Tissues were incubated with primary antibodies diluted in PBT at 4ºC. Three washes were performed the next day over several hours before nervous systems were incubated overnight at 4ºC in secondary antibodies also diluted in PBT. Tissues were then washed three times over the course of several hours and placed on cover slips coated in poly-lysine, dehydrated in an increasing ethanol concentration series, and cleared in a xylene series. Nervous systems were mounted onto slides using DPX mounting medium and imaged on a Leica TCS SP8 Confocal Microscope at 40X magnification. The following primary antibodies were used: rabbit anti-GFP (Invitrogen #A11122; 1:1000), and rat anti-DN-cadherin (DN-Ex#8, Developmental Studies Hybridoma Bank; 1:50). The following secondary antibodies were used: Fluorescein (FITC) conjugated donkey anti-rabbit (Jackson ImmunoResearch #711-095-152; 1:500), and AF-647 goat anti-rat (Invitrogen #A21247; 1:500).

### Optogenetics and Behavioral Assays

Optogenetics and behavioral assays was performed as previously described (Duckhorn et al. 2022). Briefly, unmated females used in optogenetic assays were raised in darkness and on food containing 0.2 mM all-trans-retinal (sigma-Aldrich #R2500) and were incubated at 25ºC and 50% humidity. Females were grouped in vials consisting of 15–20 flies for 8–12 days before testing. Flies were anesthetized on ice for 2 minutes, decapitated under low-intensity light, and given 15–20 minutes to recover before being transferred to individual behavioral chambers (diameter: 10 mm, height: 3 mm). A FLIR Blackfly S USB3, BFS-U3-31S4M-C camera with a 800 nm long-pass filter (Thorlabs, FEL0800) was used to record optogenetic videos in SpinView. Chambers were placed on top of an LED panel with continuous infrared (850 nm) light and recurring photoactivating red (635 nm) light using an Arduino script. To measure change in abdominal length before and during vaginal plate opening or ovipositor extrusion, a ruler (cm/mm) was included in the frame to set the scale. The change in abdominal length was calculated as the difference in the abdominal length from the base of the scutellum to the tip of the abdomen before and during photoactivation. Behavior indices were measured by calculating the average fraction of time spent performing the behavior during the first three 15-second lights-on periods and the first three 45-second light-off periods. For courtship assays, unmated females and males were collected under CO_2_ and aged for 7–10 days in a 12-hour light/dark cycle and incubated at 25ºC and 50% humidity. Unmated females were group-housed in vials consisting of 15–20 flies, and *Canton S* males were individually housed. Courtship assays were done within the first two hours of the subjective day. Unmated females and *Canton S* males were transferred to individual behavioral chambers (diameter: 10 mm, height: 3 mm) and recorded for 30 minutes using a Sony Vixia HFR700 video camera at 25ºC under white light. For experiments using mated females, unmated females were housed with males for 24 hours, anesthetized on ice for 2 minutes, and mated females were collected into a new vial and given 30 minutes to recover. Mated females and *Canton S* males were loaded to chambers and recorded as described above. Vaginal plate opening and ovipositor extrusion frequency was measured as the total number of times a female performed the behavior in a 6-min period of active male courtship. Egg laying was measured by allowing females to mate with males before transferring them to individual vials for 24 hours. The total number of eggs laid in 24 hours by each female was then counted.

## Results and Discussion

The *dsf* gene is expressed in ∼0.5% of neurons in each hemisphere of the adult central nervous system (CNS), organized in ∼11 sex-shared groups of cells in the brain and nerve cord (Figure 1A) (Duckhorn et al. 2022). One of these groups, rAbd, contain the DDAG neurons, consisting of eleven cells in females that contribute to several reproductive behaviors (Duckhorn et al. 2022). We previously showed that the DDAG neurons fall into five anatomical subtypes, A–E, and that the DDAG_C and DDAG_D subtypes regulate a specific mating behavior in mated females, *i*.*e*., ovipositor extrusion (Diamandi et al. 2024). To determine how other DDAG subtypes and other *dsf*+ neuronal groups contribute to behavior, we sought to develop reagents that provide genetic access to subsets of *dsf*-expressing neurons.

**Figure 1.**
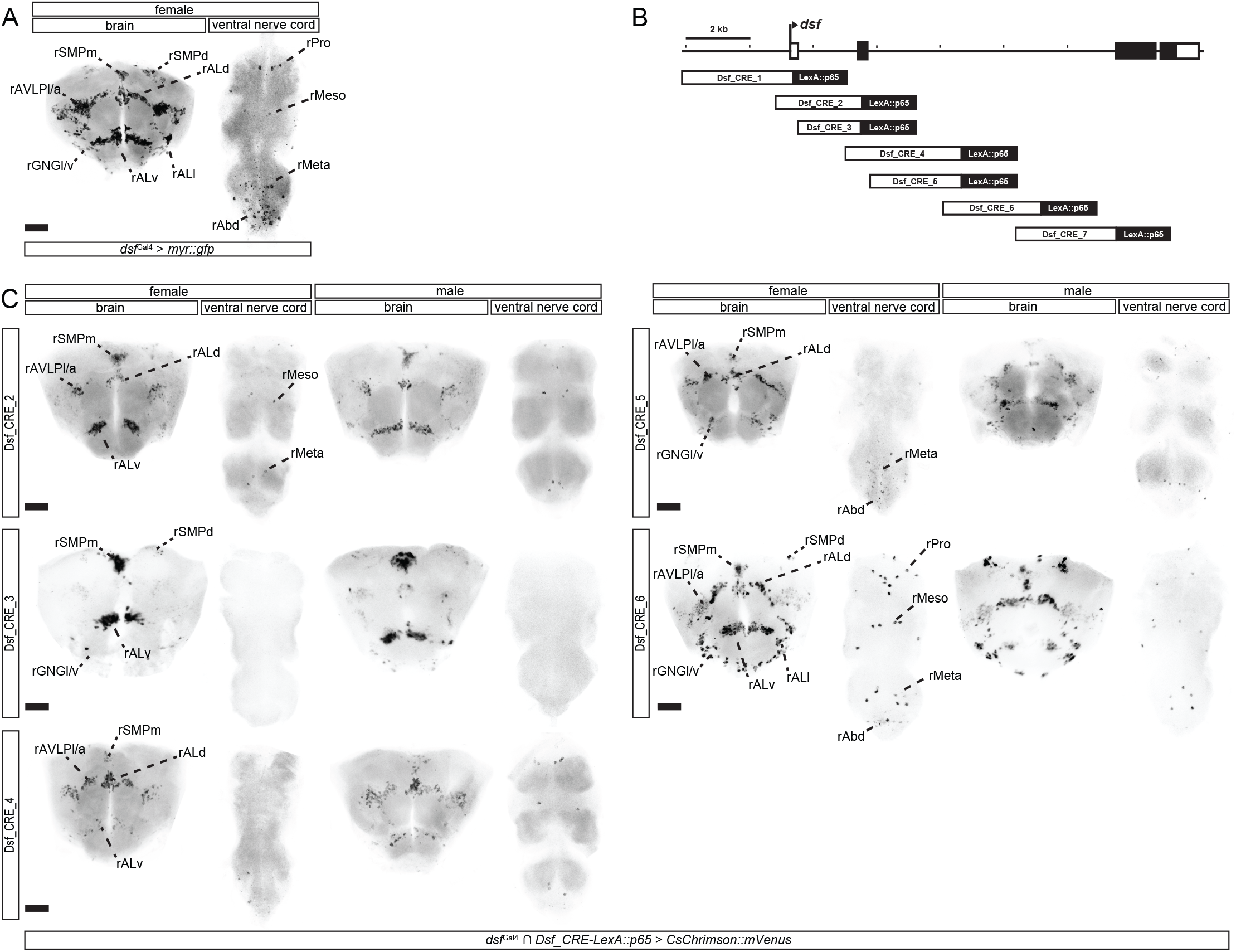
*cis*-regulatory fragments from the *dsf* gene label subsets of *dsf*-expressing neurons in the adult CNS. (A) Confocal images of the adult brain and ventral nerve cord from a *dsf*^Gal4^ > *UAS-myr::gfp* female showing expression of GFP mRNA (black) labeled by *in situ* Hybridization Chain Reaction. *Dsf*^Gal4^ is expressed in ∼11 groups of neurons in the adult brain and ventral nerve cord. *Dsf*-expressing neuronal groups were categorized according to standardized nomenclatures (Duckhorn et al. 2022). (B) An illustration showing the location of regulatory fragments from the *dsf* gene that were analyzed in this study. (C) The intersection of *dsf*^Gal4^ and most *Dsf_CRE-LexA::p65* transgene targets various subsets of *dsf*-expressing neurons in the brain and ventral nerve cord of adult females and males. Fragments 1 and 7 failed to generate any clear HCR signal and were thus not included here. Fragment 1 did, however, label a single rALv neuron by immunohistochemistry (Supplementary Figure 1). GFP mRNA is shown in black. Scale bar = 50 μm.

The noncoding regions of pleiotropic genes in metazoans often contain multiple, modular transcriptional *cis*-regulatory elements (CREs), *i*.*e*., enhancers, that direct gene expression in distinct tissues or cells. We systematically surveyed the *dsf* gene for enhancers that drive reporter expression in subsets of *dsf*-expressing cells in the CNS. We analyzed *dsf* using seven partially overlapping 2–3-kb fragments (designated as Dsf_CRE_1–7) that cover upstream and intronic regions of the locus (Figure 1B). Transgenic fly stocks were generated carrying each of the seven fragments attached to a heterologous promoter and the coding sequence for the transcriptional activator, *LexA::p65*.

To determine if these fragments drive *LexA::p65* expression in *dsf*-expressing neurons, we genetically intersected each *Dsf_CRE-LexA::p65* transgene with our *dsf*^Gal4^ allele, which faithfully recapitulates the endogenous expression of *dsf* (Duckhorn et al. 2022). In this system, LexA::p65, expressed in cells targeted by the regulatory fragment, drives the expression of a LexAop-regulated Flp recombinase, which excises a transcriptional stop cassette from a UAS-regulated *myr::gfp* transgene. Expression of GFP is then driven by *dsf*^Gal4^. We probed GFP mRNA expression in the adult CNS by *in situ* Hybridization Chain Reaction (HCR) (Figure 1C), and protein expression in the adult and larval CNS by immunohistochemistry (Supplementary Figure 1, 2). Much of *dsf*’s transcriptional *cis*-regulatory content maps to the 5’ portion of its large third intron. Most regulatory fragments in our analysis were active in subsets of *dsf*-expressing neurons in the CNS. All eleven groups of *dsf*-expressing neurons in the brain and nerve cord were labeled by one or more enhancers, and some enhancers targeted subsets of neuronal groups or subsets of cells within specific neuronal groups or both. Of the seven fragments, four (Dsf_CRE_2, 4, 5, and 6) had activity in subsets of female-specific DDAG neurons (Supplementary Figure 1). We focused our attention on Dsf_CRE_4, as this fragment labeled 3–4 female-specific DDAG neurons in each hemisphere of the abdominal ganglion and an otherwise sparse number of *dsf*-expressing neurons in the nerve cord.

Unmated *Drosophila* females indicate their receptivity to courting males with a behavior called “vaginal plate opening,” or VPO, in which they open their vaginal plates and partially expose their ovipositor (Wang et al. 2021). Mated females, however, reject courting males by fully extruding their ovipositor, a behavior called “ovipositor extrusion,” or OE, which may block copulation or male courtship drive (Wang et al. 2020). The length of the abdomen increases during both behaviors, but the change in abdominal length is greater during an OE than during a VPO (Wang et al. 2020; Duckhorn et al. 2022). We previously showed that optogenetic activation of all DDAG neurons using the mVenus-tagged, red-light-gated cation channelrhodopsin, *CsChrimson::mVenus* (Klapoetke et al. 2014), caused unmated females to open their vaginal plates and mated females to extrude their ovipositors (Duckhorn et al. 2022). Upon photoactivation, *dsf*^Gal4^ ∩*Dsf_CRE_4-LexA::p65* > *CsChrimson::mVenus* females opened their vaginal plates regardless of their mating status (Figure 2A–C; Supplementary Video 1, 2). VPO was penetrant and occurred largely during the photoactivation period (Figure 2B).

**Figure 2.**
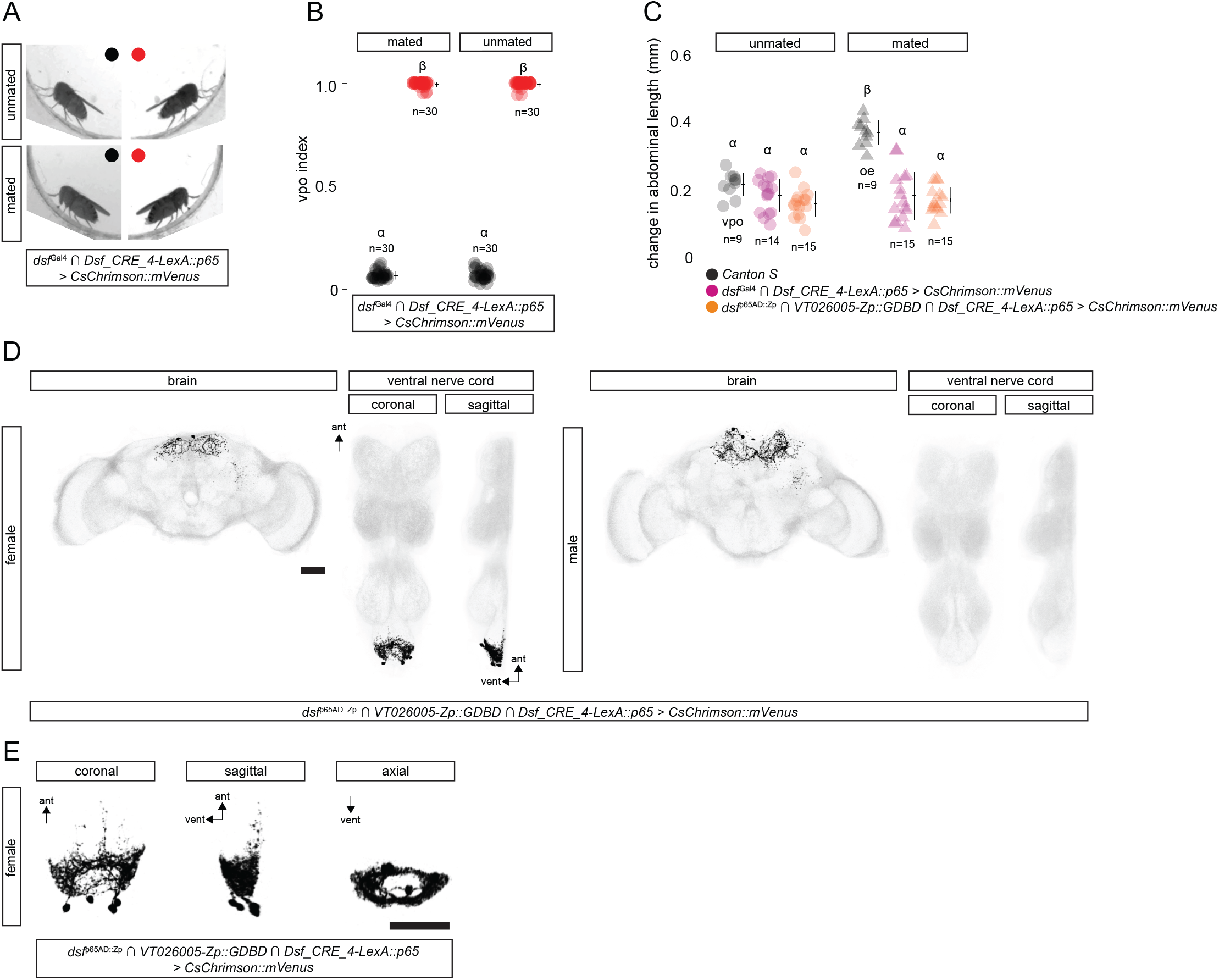
Optogenetic activation of the DDAB_B neurons opens the vaginal plates of adult females. (A) Still-frames of decapitated *dsf*^Gal4^ ∩*Dsf_CRE_4-LexA::p65* > *CsChrimson::mVenus* unmated (top) and mated (bottom) females before (left) and during (right) a bout of red light (10.5 mW/mm^2^). (B) Average fraction of time females open their vaginal plates during (red) and between (black) three 15-s photoactivation bouts in a row, *i*.*e*., VPO index. Lights are off for 45 s between bouts. It takes a few seconds for females to retract the ovipositor once the lights turn off, so indices are slightly above zero during lights off. n = number of females. (C) Photoactivation of DDAG_B neurons induces VPO in unmated and mated females. The change in abdominal length when an unmated (black circles) or mated (black triangles) *Canton S* female performs VPO or OE, respectively, during courtship is shown. Photoactivation of the DDAG_B neurons in *dsf*^Gal4^ ∩*Dsf_CRE_4-LexA::p65* > *CsChrimson::mVenus* (magenta) and *Dsf_CRE_4-LexA::p65* ∩*dsf*^p65AD::Zp^ ∩*VT026005*-*Zp::GDBD* > *CsChrimson::mVenus* (orange) unmated (circles) and mated (triangles) females induces a similar change in abdominal length to *Canton S* females performing a VPO. n = number of females. (D) Confocal images of a *Dsf_CRE_4-LexA::p65* ∩*dsf*^p65AD::Zp^ ∩*VT026005*-*Zp::GDBD* > *CsChrimson::mVenus* female and male CNS. GFP-expressing neurons and DNCad (neuropil) are shown in black and light gray, respectively. Scale bar = 50 ?m. (E) Anatomy of the DDAG_B neurons from a *Dsf_CRE_4-LexA::p65* ∩*dsf*^p65AD::Zp^ ∩*VT026005*-*Zp::GDBD* > *CsChrimson::mVenus* female. Scale bar = 50 μm.

The female-specific neurons labeled by *dsf*^Gal4^ ∩*Dsf_CRE_4-LexA::p65* appeared to correspond to a single anatomical subtype we previously identified called the DDAG_B neurons (Diamandi et al. 2024). The DDAG_B neurons are female-specific interneurons that arborize locally in the abdominal neuropil, forming a hollow circle in an axial view (Figure 2E). We previously developed a split-gal4 driver, *dsf*^p65AD::Zp^ ∩*VT026005*-*Zp::GDBD*, that labeled the DDAG_C, DDAG_D, and DDAG_B neurons in adult females (Diamandi et al. 2024). To confirm that the DDAG neurons labeled by Dsf_CRE_4 are the DDAG_B neurons, we genetically intersected *Dsf_CRE_4-LexA::p65* with *dsf*^p65AD::Zp^ ∩*VT026005*-*Zp::GDBD*. Indeed, this three-way intersection specifically labeled 3–4 female-specific DDAG_B neurons and no other neurons in the nerve cord (Figure 2D, E). Photoactivation of *Dsf_CRE_4-LexA::p65* ∩*dsf*^p65AD::Zp^ ∩*VT026005*-*Zp::GDBD* > *CsChrimson::mVenus* females evoked VPO behaviors (Figure 2B, C; Supplementary Video 3) that were qualitatively and quantitatively similar to what we observed when we photoactivated *dsf*^Gal4^ ∩*Dsf_CRE_4-LexA::p65* > *CsChrimson::mVenus* females.

To determine if activity of the DDAG_B neurons labeled by *Dsf_CRE_4-LexA::p65* ∩*dsf*^p65AD::Zp^ ∩*VT026005*-*Zp::GDBD* is required for VPO behavior in unmated females during courtship, we used the three-way intersection to drive the expression of a GFP-tagged inwardly rectifying K+ channel, Kir2.1::gfp (Baines et al. 2001) (Figure 3A). We observed no effect on female courtship behaviors from the expression of *Kir2*.*1::gfp* in the DDAG_B neurons. Unmated and mated females expressing *Kir2*.*1::gfp* in the DDAG_B neurons copulated with males at a rate that was similar to control females (Figure 3B, C); they opened their vaginal plates and extruded their ovipositors during courtship at a frequency comparable to controls (Figure 3D, E); and mated *Dsf_CRE_4-LexA::p65* ∩*dsf*^p65AD::Zp^ ∩*VT026005*-*Zp::GDBD* > *Kir2*.*1::gfp* females laid a number of eggs similar to controls (Figure 3F). These data suggest the involvement of additional neural circuit elements for VPO in the abdominal ganglion that may compensate for the absence of DDAG_B activity. Alternatively, additional DDAG_B neurons may exist that were not targeted by our labeling strategy.

**Figure 3.**
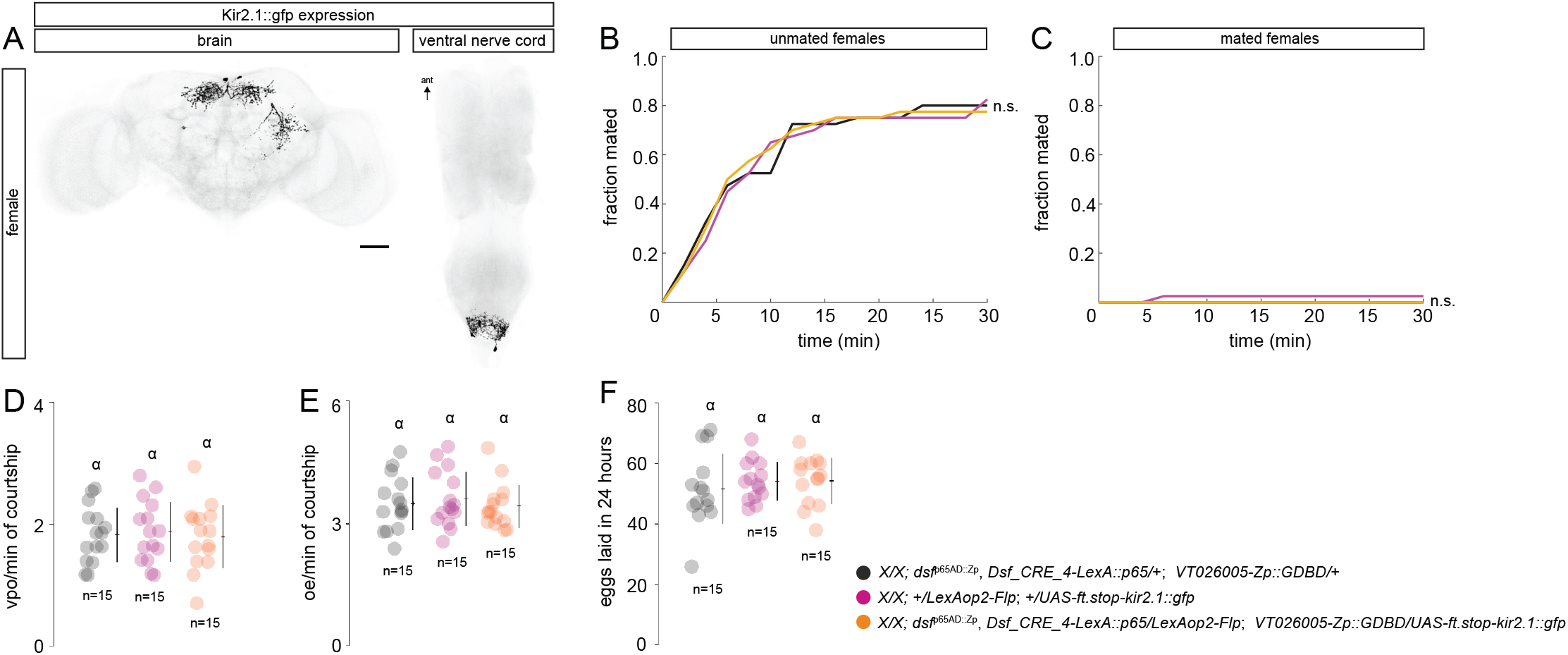
Expression of Kir2.1::GFP in the DDAG_B neurons has no effect on female courtship behaviors. (A) Confocal image of a *Dsf_CRE_4-LexA::p65* ∩*dsf*^p65AD::Zp^ ∩*VT026005*-*Zp::GDBD* > *kir2*.*1::gfp* female brain and ventral nerve cord. Kir2.1::GFP-expressing neurons are in black. (B, C) Fraction of unmated (B) and mated (C) females that mated with naive *Canton S* males in 30 min. Log rank test for significance. n.s. = not significant. n = 40 females for each genotype. (D) Number of VPOs an unmated female performed per minute of active courtship. (E) Number of OEs a mated female performed per minute of active courtship. (F) Number of eggs laid 24-h post-mating. (D–F) Individual points, mean, and SD. n = number of females. A one-way ANOVA Tukey-Kramer multiple comparison test measured significance (p < 0.05). Same letter indicates no significant difference.

The DDAG neurons are an anatomically diverse population consisting of at least five subtypes that collectively influence multiple female mating behaviors (Duckhorn et al. 2022; Diamandi et al. 2024). We previously showed that the DDAG_C and DDAG_D neurons regulate OE in mated females (Diamandi et al. 2024). Here, we found that the DDAG_B neurons likely contribute to VPO in unmated females. Our results suggest that the motor circuits for VPO and OE in the abdominal ganglion of females may include neurons with functions specific to each behavior. The EM connectome data of the female and male ventral nerve cord (Marin et al. 2024; Stürner et al. 2025) will provide important insights into the position of these neurons in the abdominal circuits that modulate female abdominal behaviors during courtship.

The DDAG neurons are all recycled from a population of *dsf*-expressing interneurons in the larval abdominal ganglion (Diamandi et al. 2024). The DDAG_C and DDAG_D neurons are segmental homologs of a single, sex-shared larval interneuron type called A26g (Diamandi et al. 2024), whose function is currently unknown. During early metamorphosis, A26g neurons at abdominal segments A5 and A6 acquire *dsx* expression, undergo programmed cell death in males, and get repurposed in females for functions in ovipositor extrusion (Diamandi et al. 2024). What about the DDAG_B neurons? In future studies, it will be fascinating to identify the larval counterparts of the DDAG_B neurons and determine how they contribute to larval life.

## Acknowledgements

We thank J. Cande and D. Stern (HHMI) for contributing the *Dsf_CRE-LexA::p65* transgenes; E. Preger-Ben Noon for helpful comments on the manuscript; and A. McStravog for administrative assistance.

T.R.S. is supported by the National Science Foundation (IOS-1845673, IOS-2445300).

## Figure legends

**Supplementary Figure 1. *cis*-regulatory fragments from the *dsf* gene label subsets of *dsf*-expressing neurons in the adult CNS**. The intersection of *dsf*^Gal4^ and each *Dsf_CRE-LexA::p65* transgene targets various subsets of *dsf*-expressing neurons in the brain and ventral nerve cord of adult females and males. GFP-expressing neurons and DNCad (neuropil) are shown in black and light gray, respectively. Scale bar = 50 μm.

**Supplementary Figure 2. *cis*-regulatory fragments from the *dsf* gene label subsets of *dsf*-expressing neurons in the larval CNS**. The intersection of *dsf*^Gal4^ and each *Dsf_CRE-LexA::p65* transgene targets various subsets of *dsf*-expressing neurons in the brain and ventral nerve cord of larval females and males. GFP-expressing neurons and DNCad (neuropil) are shown in black and light gray, respectively. Scale bar = 50 μm.

**Supplementary Video 1. Optogenetic activation of *dsf***^**Gal4**^ ∩***Dsf_CRE_4-LexA::p65* > *CsChrimson::mVenus* unmated females**. Upon photoactivation, a *dsf*^Gal4^ ∩*Dsf_CRE_4-LexA::p65* > *CsChrimson::mVenus* unmated female opens her vaginal plates.

**Supplementary Video 2. Optogenetic activation of *dsf***^**Gal4**^ ∩***Dsf_CRE_4-LexA::p65* > *CsChrimson::mVenus* mated females**. Upon photoactivation, a *dsf*^Gal4^ ∩*Dsf_CRE_4-LexA::p65* > *CsChrimson::mVenus* mated female opens her vaginal plates, similar to an unmated female.

**Supplementary Video 3. Optogenetic activation of *Dsf_CRE_4-LexA::p65* ∩*dsf***^**p65AD::Zp**^ **∩*VT026005*-*Zp::GDBD* > *CsChrimson::mVenus* unmated females**. Upon photoactivation, a *Dsf_CRE_4-LexA::p65* ∩*dsf*^p65AD::Zp^ ∩*VT026005*-*Zp::GDBD* > *CsChrimson::mVenus* unmated female opens her vaginal plates.

## Notes

### Competing Interest Statement

The authors have declared no competing interest.

### Summary of Updates

The first submission of the manuscript to bioRxiv occurred through the journal we submitted the paper to and did not include the Supplementary Data. In the updated version includes Supplementary Figures 1, 2 and Videos 1 to 3.

