## Supplementary Figure 2 for "*Cis*-regulatory fragments from the *dissatisfaction* gene identify novel mating behavior neurons in female *Drosophila*"

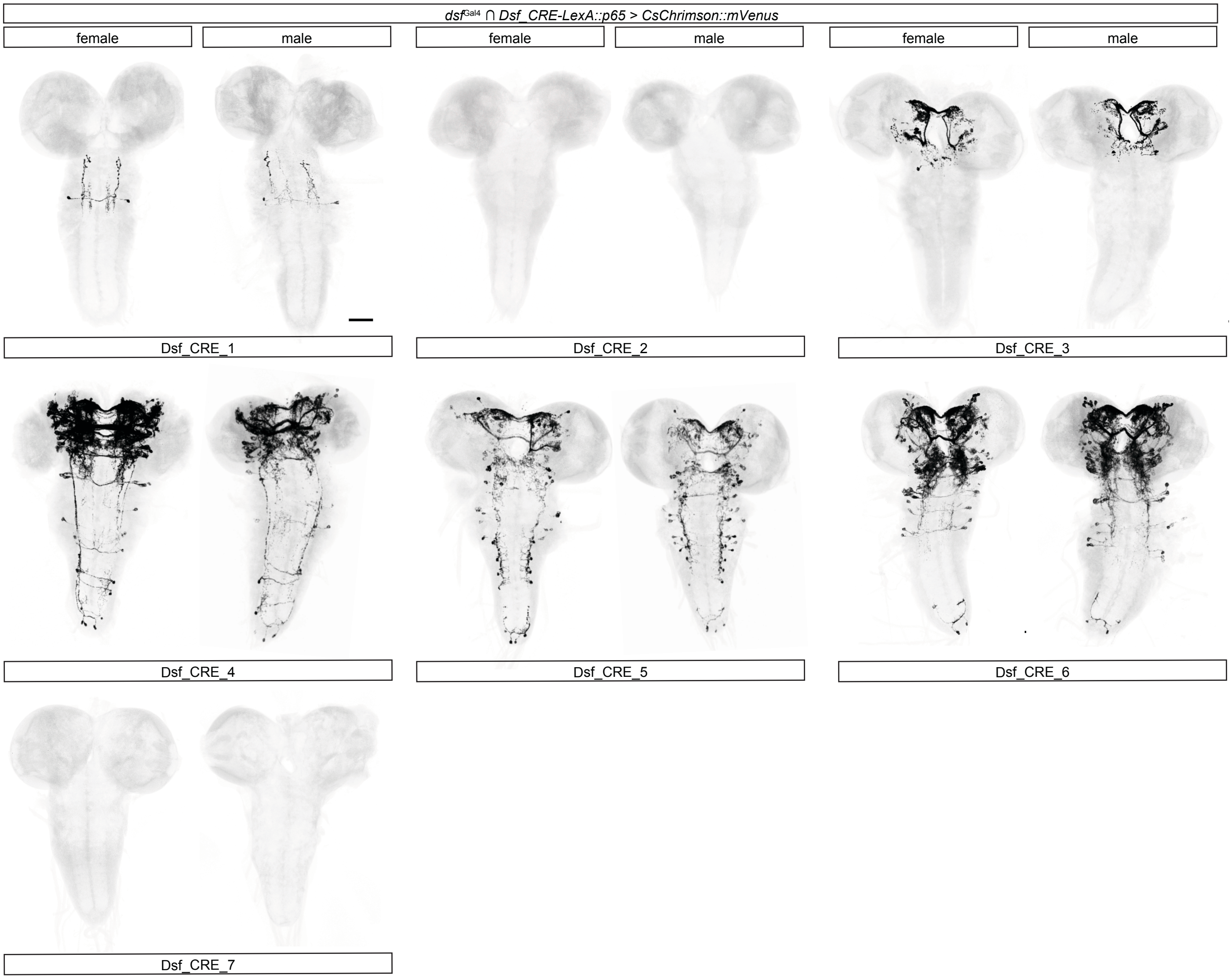


**Supplementary Figure 2**. cis-regulatory fragments from the dsf gene label subsets of dsf-expressing neurons in the larval CNS. The intersection of dsfGal4 and each Dsf_CRE-LexA::p65 transgene targets various subsets of dsf-expressing neurons in the brain and ventral nerve cord of larval females and males. GFP-expressing neurons and DNCad (neuropil) are shown in black and light gray, respectively. Scale bar = 50 m.
